# Hearts in sync on a roller coaster: heart-rate responses, synchrony and psychological change in 439 amusement-park riders

**DOI:** 10.64898/2026.11.19.767997

**Authors:** Daniel Berglind, Charlotte Wilén, Mehedi Hasan, Greta Wilsson, Micael Dahlén

**Author notes:** Corresponding author: Daniel Berglind.

## Abstract

**Background:** Shared intense experiences are thought to bring people closer, and synchronized physiology is a proposed reason. We examined heart rate and synchrony across ride phases, riders’ response profiles, and whether these relate to changes in happiness, stress and closeness.

**Methods:** In this exploratory field study we recorded heart rate in 439 riders (418 with usable data) on 19 trains of the Balder roller coaster, Gothenburg, Sweden, and measured happiness, stress and closeness (0-10) just before and after the ride. Real pairs were compared with surrogate pairs from different trains, aligned on ride start, to separate coupling between riders from the ride’s own effect. Profiles came from k-means clustering; analyses used linear mixed models.

**Results:** Heart rate rose by 15.2 beats per minute (95% CI 14.2 to 16.3) during the two-minute ride and was back at baseline within nine minutes. Heart-rate curves did not form clearly distinct types (two profiles, silhouette 0.29): riders calm in the queue rose more and stayed elevated, while riders already aroused rose less. Pairs in which both riders came with companions gained more synchrony during the ride than the surrogate benchmark (Fisher z 0.096, 95% CI 0.050 to 0.142); for pairs including a solo rider the boost was smaller and uncertain. This coupling peaked just after the ride and was gone five minutes later. Riders reported being happier (+1.3 points, Cohen’s d = 0.87) and closer to the other riders on the train (+2.2 points, d = 0.70). Neither phase-specific heart-rate responses, response profile nor synchrony was associated with these changes.

**Conclusions:** A short shared thrill coupled riders’ heart rates beyond what the ride alone explains, and riders felt happier and closer to others on the train afterwards, but the two ran in parallel. Without a non-riding comparison, the psychological changes cannot be attributed to the ride itself.

## Introduction

People often describe intense experiences shared with others as among the most bonding moments of their social lives, and experimental work shows that sharing an experience amplifies how it feels [1,2]. A popular explanation is interpersonal physiological synchrony: the tendency of two people’s bodily arousal to rise and fall together during a joint activity [3,4]. Synchrony has been observed from parent-infant play to collective rituals [4,5], and recent field studies link it to group performance and everyday social engagement [6,7]. It is therefore often read as a bodily signature of social connection.

Two problems make that reading hard to test outside the laboratory. First, two people who go through the same event at the same time will show similar physiology simply because the event drives both of them. A claim that synchrony is *interpersonal* needs a benchmark for how much similarity the shared event alone produces [8]. Second, people do not respond to the same event in the same way, and when a response happens may matter as much as how large it is. Visitors to an extreme haunted attraction reported better mood afterwards than before [9], and in a haunted-house field study enjoyment tracked moment-to-moment heart-rate fluctuations rather than overall fear [10]. Averaging across everyone, or across the whole experience, can therefore hide distinct response profiles, and it can hide associations that only exist in one part of the experience, such as the anticipation before it starts or the recovery afterwards.

A roller coaster is a well-controlled natural laboratory for both problems. The ride is intense, short and mechanically identical from one train to the next; a full train is exposed at once; seats are assigned and known; and riders arrive both with friends and family and alone. Riders on *other* trains of the same ride provide the benchmark: they share the ride’s timeline but never shared a vehicle.

We recorded heart rate and brief self-reports from 439 riders of the Balder roller coaster at Liseberg amusement park in Gothenburg, Sweden. We asked three questions. (1) How does heart rate change across the phases of the ride, and do riders fall into distinct response profiles? (2) Do riders’ heart rates synchronise beyond what the shared ride explains, in which phases, and does this depend on whether they came with companions or sat close to each other? (3) Are phase-specific heart-rate responses, response profiles or synchrony related to the changes in happiness, stress and closeness that riders report? The study was exploratory and observational: it describes what happened around a shared ride rather than testing what the ride caused.

## Materials and methods

### Design and setting

This was a prospective observational field study with repeated measures within each rider, reported following the STROBE guidelines [11] (S1 Checklist). Riders of Balder, a wooden roller coaster at Liseberg amusement park in Gothenburg, Sweden, were recruited on site between 14 July and 21 August 2026. A Balder train seats 30 riders in five cars of three rows with two seats each. Riders did not choose their seats: they were placed according to their participant ID. Riders who came with others were therefore not necessarily seated next to them, and could end up beside people they had not come with. Data were collected in 14 measurement groups that filled 19 trains (“dispatches”). Nine groups fitted in one train; five did not, and the riders left over took the next train: 7 riders in Group 8 and 2-3 riders in each of Groups 9, 11, 12 and 13 (S3 Table). Because riders on the same train share the stimulus, the train is the unit for everything shared between riders.

### Heart rate

Each rider wore a Polar Ignite 3 wrist-worn optical heart-rate monitor during the visit, recording at 1 Hz. One numbered device was assigned per seat.

### Self-report

**I**mmediately before and immediately after the ride, riders rated on 0-10 scales how happy and how stressed they felt, how close they felt to the people they came with, and how close they felt to the other riders. After the ride they also rated how intense the ride felt, how safe they felt, whether they wanted to ride again, and any discomfort (S1 Table). Riders who reported riding without companions are called solo riders. The upper anchor of the two closeness items was “very close” before the ride and “extremely close” after it (S1 Table); scores were analysed as recorded. The survey did not ask whether riders already knew the other people on the train.

### Session context

For each train a researcher logged the departure time, queue time, weather, time of day and park occupancy (S3 Table).

### Ride phases

All riders on a train start at the same moment. We therefore detected the ride start (onset) for each train from the group-average heart-rate curve, as the steepest sustained rise near the researcher-logged departure, and checked it against that log (S2 Fig). For the four second trains with only 2-3 riders, the group average rests on very few curves and the detected onset is uncertain (S3 Table). These 11 riders were kept in the main analyses; leaving them out changed none of the results (S2 Table). Each rider’s recording was then cut into four phases: baseline (the 60 s before onset), ride (0-120 s), early recovery (120-240 s) and late recovery (240-540 s). Within the first minutes we also used each rider’s own timing: calm queueing (first 30 s of baseline), anticipation (the last 30 s before onset), the rider’s own peak (±15 s around their maximum during the ride) and relief (the 30 s after that peak).

### Heart-rate response profiles

To describe how riders differ, we grouped individual heart-rate curves by shape. For each rider we expressed heart rate relative to their own baseline, averaged it in 10 s steps from 60 s before to 540 s after onset (riders with at least 80% of steps covered; gaps interpolated), and clustered these curves with k-means. We compared solutions with two to six profiles and chose the one with the best average silhouette width [12], provided no profile held fewer than 8% of riders. Stability was checked by re-clustering 100 bootstrap samples and comparing the result with the original solution [13]. Profiles are numbered from the strongest to the weakest peak response.

### Heart-rate synchrony

Synchrony between two riders was the correlation between their baseline-centred heart-rate curves within a phase, Fisher z-transformed, requiring at least 30 s of overlapping data. Real pairs were two riders on the same train, classed as companion pairs (both reported coming with companions) or stranger pairs (at least one rode alone). A companion pair is therefore two riders who each came with someone, but not necessarily with each other or seated next to each other. The comparison of companion and stranger pairs is thus a comparison between riders who came with someone and riders who came alone, not between people who know each other and people who do not. The surrogate benchmark paired riders from different trains, aligned on time since their own onset [8]. Such pairs share the ride’s timing but never shared a vehicle, so any synchrony in them comes from the ride alone. The quantity of interest is the gap between real pairs and the benchmark. As a robustness check we also correlated second-to-second changes instead of levels (first-differenced synchrony), which removes slow shared trends. Seat proximity was classed as side by side, same car but different row, or different car.

### Statistical analysis

We used linear mixed models throughout. Heart rate was modelled by phase with a random intercept per rider, adjusting for queue time, temperature and time of day. Synchrony was modelled with phase, pair type and their interaction, with a random intercept per pair and the surrogate benchmark as reference. In the primary model (baseline and ride), the interaction is the ride-induced boost beyond the benchmark. The extended model added the two recovery phases. Contrasts were tested with Wald tests, and the key companion-versus-stranger contrast was also checked with a bootstrap that resampled whole trains.

Changes in self-reports from before to after the ride were tested with within-person mixed models. To ask whether physiology is related to psychology, we related four phase-specific heart-rate responses (anticipation, ride, peak and late recovery, each scaled per standard deviation) to six outcomes (changes in happiness, stress and the two closeness ratings, and rated intensity and wish to ride again), with a random intercept per train. Response profiles were compared on rider characteristics (chi-square tests) and outcomes (likelihood-ratio tests in mixed models). Pairwise synchrony was related to the pair’s average change in closeness to companions. Within each family of tests, we controlled the false discovery rate with the Benjamini-Hochberg procedure [14]. Tests were two-sided with α = .05. Analyses were run in Python 3 (pandas, statsmodels, SciPy, scikit-learn).

The analyses were exploratory and not preregistered. The phase-by-phase synchrony, phase-specific heart-rate and response-profile analyses were added after the primary analyses, and all estimates from them are reported (S5-S7 Tables). The emphasis on happiness, stress and closeness reflects the questions of this report, not prospectively designated primary outcomes.

### Ethics statement

This study was submitted to the Swedish Ethical Review Authority (Etikprövningsmyndigheten). The Authority determined that the study did not require formal ethical review under the Swedish Ethical Review Act (Dnr 2026-04186-01). All participants provided written informed consent prior to participation, and the study was conducted in accordance with the Declaration of Helsinki.

## Results

### Participants

In total 439 riders took part (362 with companions, 73 alone). Of these, 418 (95.2%) had usable heart-rate data and 402 (91.6%) had data through the full recovery window (S1 Fig). Most were experienced Balder riders: 340 (77.4%) had ridden it more than three times and 21 (4.8%) were first-time riders (Table 1).

**Table 1.** Participant characteristics.

|  | **All riders** | **With companions** | **Riding alone** |
|---|---|---|---|
| n | 439 | 362 | 73 |
| Happiness before the ride (0-10), mean (SD) | 7.5 (1.8) | 7.6 (1.7) | 7.2 (1.9) |
| Stress before the ride (0-10), mean (SD) | 3.5 (2.7) | 3.5 (2.7) | 3.2 (2.6) |
| First ride on Balder, n (%) | 21 (4.8%) | 20 (5.5%) | 1 (1.4%) |
| Ridden 1-3 times before, n (%) | 74 (16.9%) | 60 (16.6%) | 14 (19.2%) |
| Ridden more than 3 times, n (%) | 340 (77.4%) | 282 (77.9%) | 58 (79.5%) |
| Usable heart-rate data, n (%) | 418 (95.2%) | 345 (95.3%) | 70 (95.9%) |
| Heart rate through full recovery, n (%) | 402 (91.6%) | 332 (91.7%) | 67 (91.8%) |
| Reported serious discomfort, n (%) | 3 (0.7%) | 1 (0.3%) | 2 (2.7%) |

Riders with missing companion information are included in the first column only.

### Heart rate across the ride phases

Heart rate rose from 94.8 bpm at baseline to 110.0 bpm during the ride, an increase of 15.2 bpm (95% CI 14.2 to 16.3; Fig 1A, Table 2). It was still 11.2 bpm above baseline in early recovery, and back at baseline in late recovery (+0.7 bpm, 95% CI -0.4 to 1.8; P = .192). The rise started before the train moved: in the last 30 s before departure heart rate was 4.3 bpm above calm queueing (95% CI 3.3 to 5.3). Each rider’s own peak was 26.4 bpm above calm queueing, and 30 s later heart rate had fallen by only 1.8 bpm (Fig 1B).

**Fig 1.**
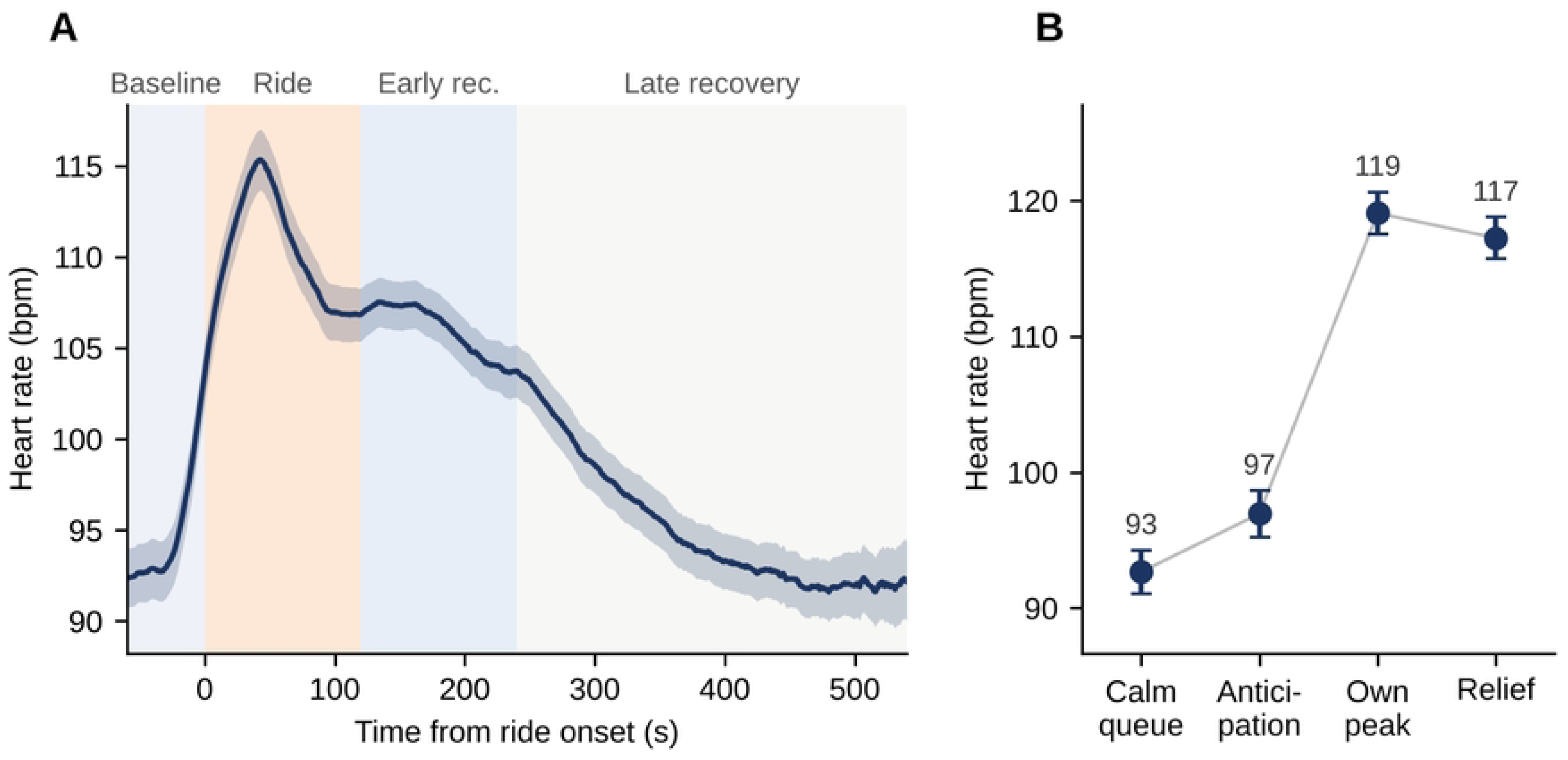
Heart rate before, during and after the ride. (A) Mean heart rate (line) with 95% confidence band across riders, by time from ride onset. Shading marks baseline, ride, early recovery and late recovery. (B) Mean heart rate (±95% CI) in four individually timed windows: calm queueing, anticipation (last 30 s before departure), the rider’s own peak and the 30 s after it. n = 418 riders (402 in late recovery).

**Table 2.** Heart rate by ride phase.

|  | **Mean HR, bpm (SD)** | **Difference, bpm (95% CI)** | **P** |
|---|---|---|---|
| Ride phases (group-timed) |  |  |  |
| Baseline (60 s before departure) | 94.8 (16.8) | Reference |  |
| Ride (0-2 min) | 110.0 (15.3) | +15.2 (14.2 to 16.3) | <.001 |
| Early recovery (2-4 min) | 106.0 (13.0) | +11.2 (10.1 to 12.2) | <.001 |
| Late recovery (4-9 min) | 95.6 (13.5) | +0.7 (-0.4 to 1.8) | .192 |
| Within the ride (individually timed) |  |  |  |
| Calm queueing (first 30 s of baseline) |  | Reference |  |
| Anticipation (last 30 s before departure) |  | +4.3 (3.3 to 5.3) | <.001 |
| Own peak (±15 s) |  | +26.4 (25.4 to 27.4) | <.001 |
| Relief (30 s after own peak) |  | +24.6 (23.6 to 25.6) | <.001 |

Differences from linear mixed models with a random intercept per rider, adjusted for queue time, temperature and time of day (ride phases) or unadjusted (within-ride windows). HR, heart rate; bpm, beats per minute.

### Heart-rate response profiles

Riders’ heart-rate curves varied a lot, but they did not fall into clearly separate types. The best clustering had 2 profiles, and its structure was weak: the silhouette width was 0.29, where values below about 0.25 indicate no real structure and 0.26-0.50 weak structure [15], and it fell steadily with more profiles (S4 Fig, S7 Table). The solution was, however, stable (bootstrap adjusted Rand index 0.84).

The two profiles mainly separated riders by how aroused they already were in the queue (Fig 2A, Table 3). The profiles cover the 385 riders with a near-complete curve. In profile 1 (n = 177, 46.0%), riders started from a calm baseline (85 bpm), peaked 31.7 bpm above it after about 66 s, and were still 10 bpm above it in late recovery. In profile 2 (n = 208, 54.0%), riders were already aroused before departure (104 bpm), peaked earlier and lower (21.7 bpm after about 45 s), and ended 9 bpm below their queue level. This is the familiar pattern that a response depends on the starting level. The profiles did not differ in companionship (82% vs. 84% with companions), first rides (5% vs. 5%), happiness before the ride, or any psychological outcome (all q ≥ 0.41; Fig 2B, Table 3).

**Fig 2.**
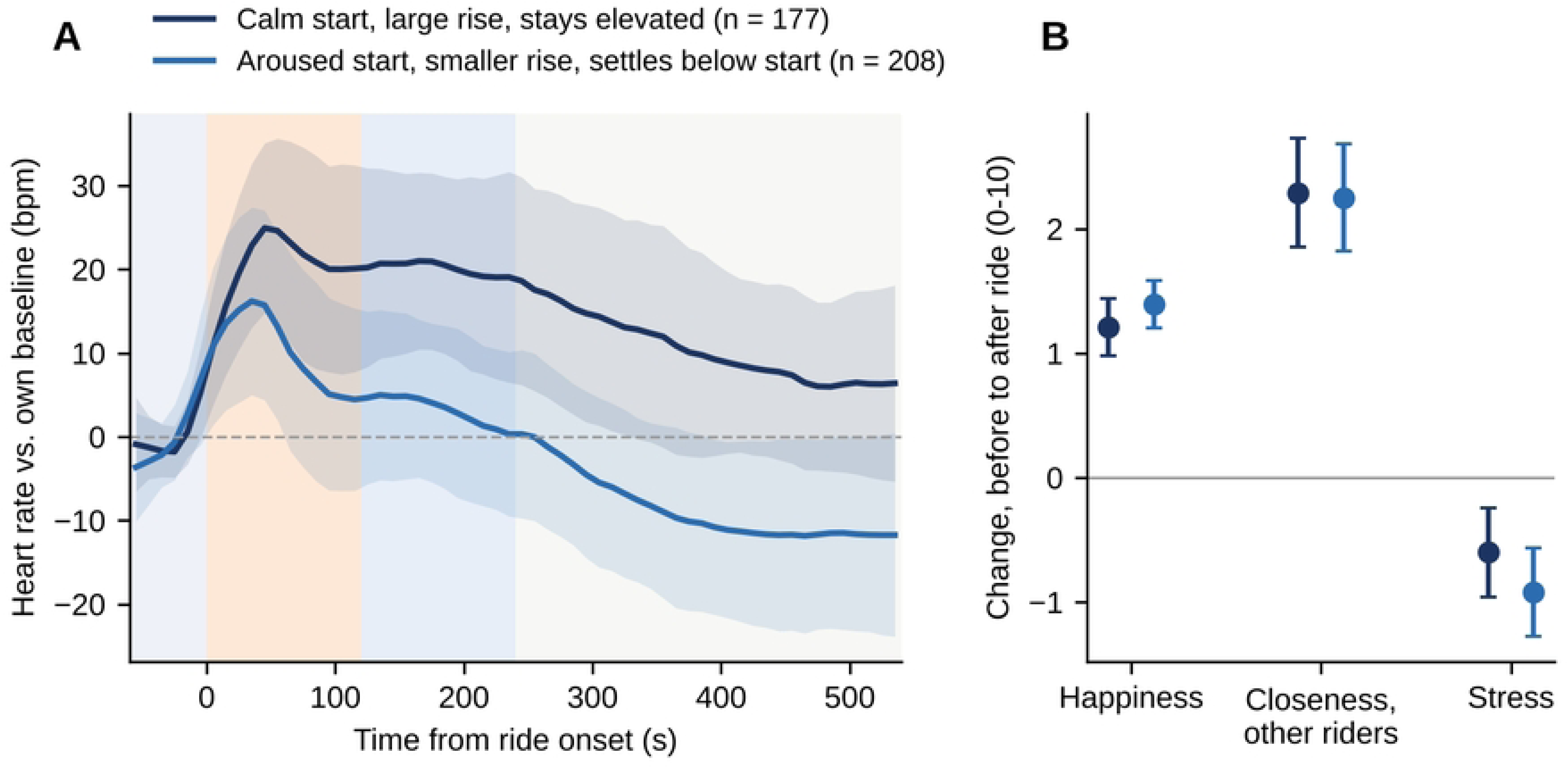
Heart-rate response profiles. (A) Mean heart rate relative to each rider’s own baseline for each profile (line) ±1 SD (band). Shading as in Fig 1. (B) Mean change (±95% CI) from before to after the ride in happiness, closeness to other riders and stress, by profile (colours as in A).

**Table 3.** Heart-rate response profiles: shape, rider characteristics and outcomes.

|  | **Profile 1** | **Profile 2** | **P** |
|---|---|---|---|
| n (%) | 177 (46.0) | 208 (54.0) |  |
| Anticipation rise, bpm | 1.3 | 2.9 |  |
| Peak rise, bpm, mean (SD) | 31.7 (9.9) | 21.7 (10.0) |  |
| Time to peak, s, mean (SD) | 66 (30) | 45 (30) |  |
| Late recovery vs. baseline, bpm | 10.5 | -8.6 |  |
| Riding with companions (%) | 82 | 84 | .745 |
| First ride on Balder (%) | 5 | 5 | .923 |
| Baseline heart rate (bpm) | 85.0 (12.2) | 103.9 (15.6) | <.001 |
| Happiness before the ride (0-10) | 7.50 (1.89) | 7.51 (1.65) | 1.000 |
| Change in happiness | 1.21 (1.54) | 1.40 (1.40) | .221 |
| Change in stress | -0.60 (2.41) | -0.92 (2.60) | .218 |
| Change in closeness to other riders | 2.29 (2.93) | 2.25 (3.13) | .886 |
| Change in closeness to companions | 0.08 (1.52) | 0.28 (1.54) | .239 |
| Rated intensity (0-10) | 7.90 (1.51) | 8.12 (1.47) | .151 |
| Wants to ride again (0-10) | 8.91 (2.08) | 8.98 (2.01) | .835 |

Values are mean (SD) or %, unless stated. P values from chi-square tests (percentages) or likelihood-ratio tests in mixed models with a random intercept per train (means). Outcomes are changes from before to after the ride unless stated.

### Synchrony between riders across the ride phases

Average synchrony was low to moderate for all pair types (companion pairs n = 3812, stranger pairs n = 1670, surrogate pairs n = 3000). It was higher before the ride than during it for every pair type, including the benchmark (benchmark change -0.136, 95% CI -0.170 to -0.101; Fig 3). This reflects the slow, shared rise in the queue, which makes any two curves look alike, while peaks during the ride come at different moments for different riders. The benchmark captures exactly this, so the gap between real pairs and the benchmark is what matters.

**Fig 3.**
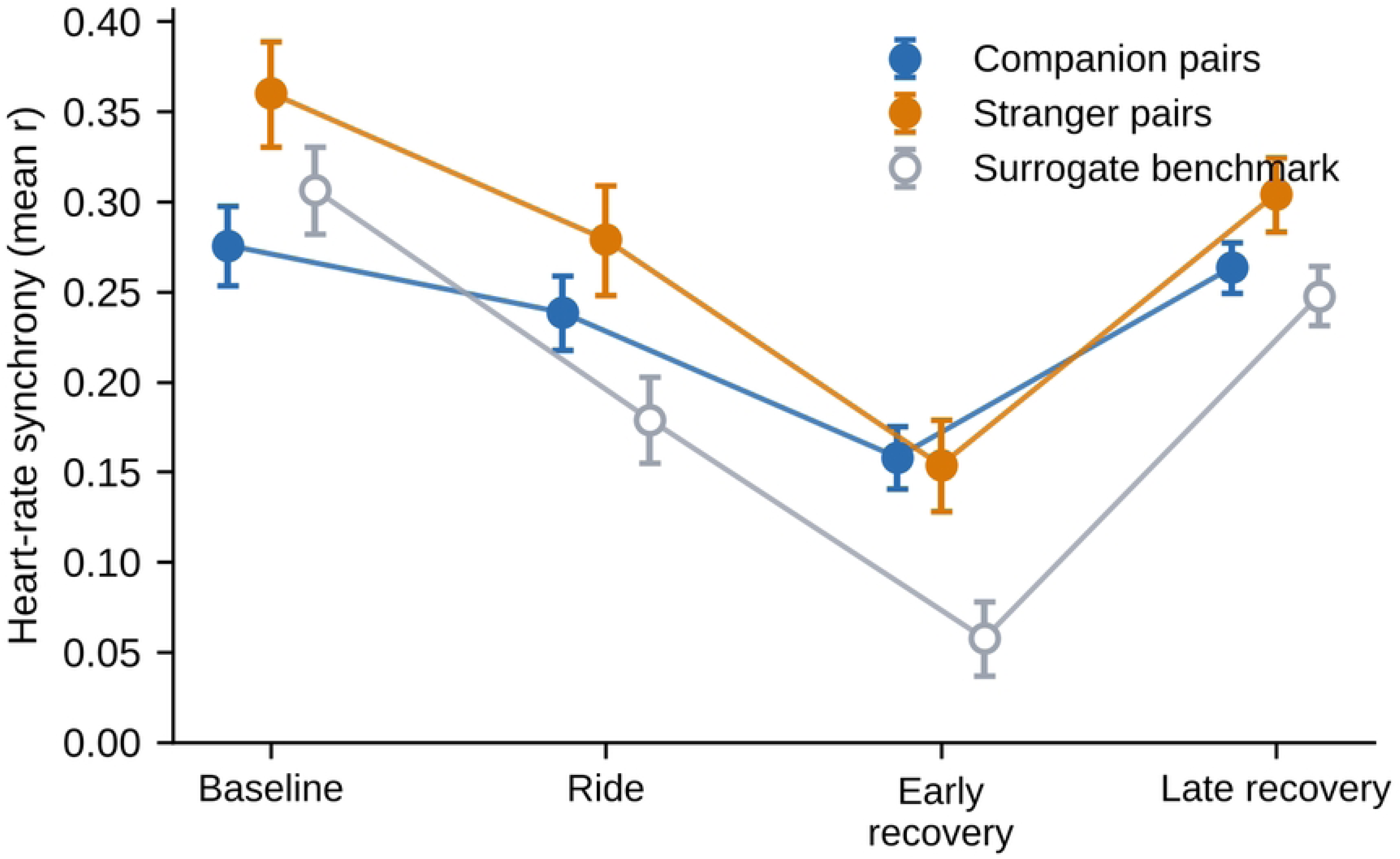
Heart-rate synchrony between riders in each ride phase. Mean synchrony (correlation, back-transformed from Fisher z; ±95% CI) for companion pairs, stranger pairs on the same train and the surrogate benchmark (riders from different trains aligned on ride onset). The benchmark shows how similar two riders’ curves are because of the ride alone; the gap between a real pair type and the benchmark is the coupling between riders.

On that measure, companion pairs gained more synchrony from baseline to ride than the benchmark (ride boost 0.096, 95% CI 0.050 to 0.142; P < .001; Table 4). Stranger pairs showed a smaller boost that was not statistically clear (0.045, 95% CI -0.013 to 0.103; P = .126). The direct comparison did not clearly favour either pair type (-0.051, 95% CI -0.106 to 0.005; P = .074); 81.0% of train-level bootstrap samples pointed towards companions, but the bootstrap interval included zero (-0.149 to 0.061). Companion pairs started out less synchronised than stranger pairs (Table 4), so a larger boost for them would not be a ceiling effect. The first-differenced check pointed the other way: stranger pairs, not companions, exceeded the benchmark (0.021, 95% CI 0.003 to 0.040; P = 0.019; companions 0.007, P = 0.407; S2 Table). We therefore treat the companion-versus-stranger difference as unresolved.

**Table 4.**
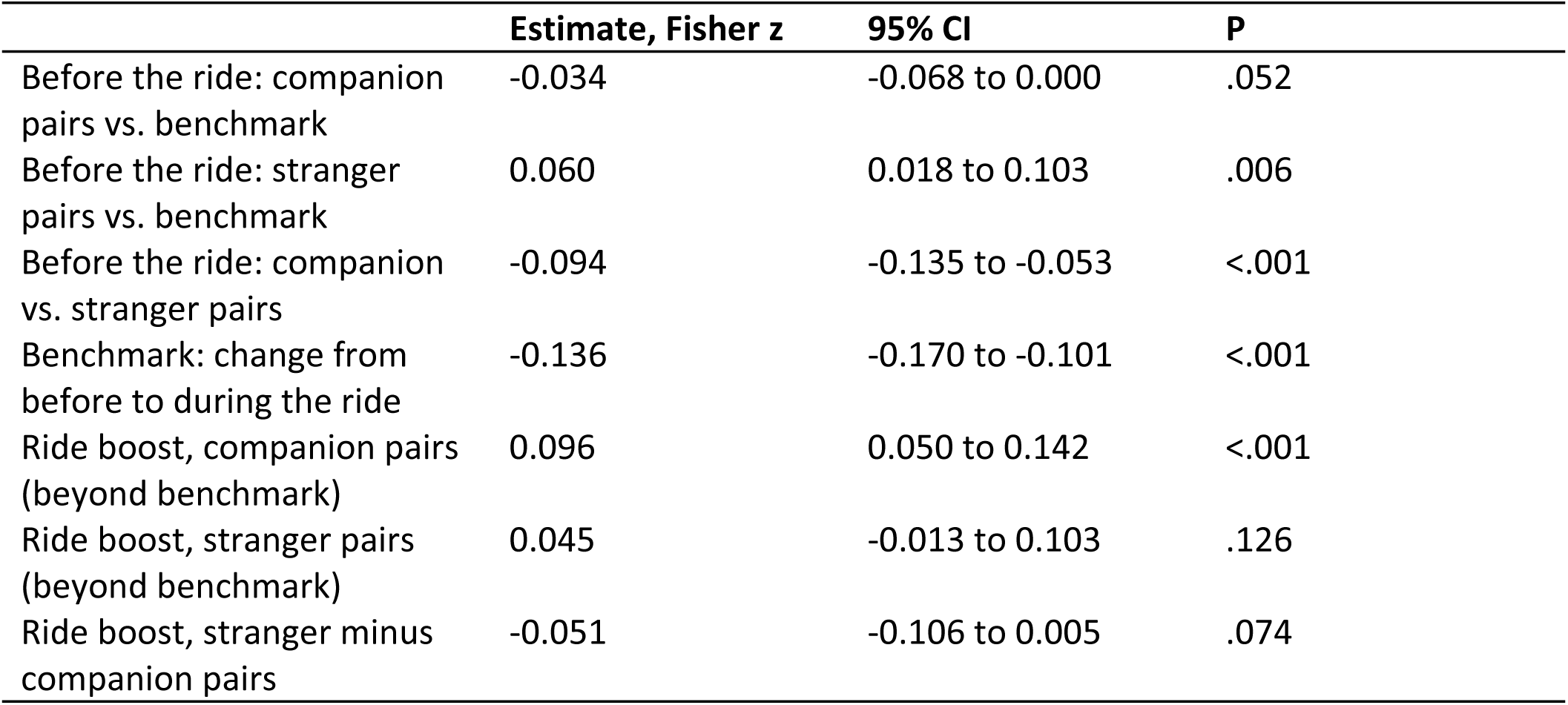
Heart-rate synchrony: baseline levels and ride-induced boosts.

|  | **Estimate, Fisher z** | **95% CI** | **P** |
|---|---|---|---|
| Before the ride: companion pairs vs. benchmark | -0.034 | -0.068 to 0.000 | .052 |
| Before the ride: stranger pairs vs. benchmark | 0.060 | 0.018 to 0.103 | .006 |
| Before the ride: companion vs. stranger pairs | -0.094 | -0.135 to -0.053 | <.001 |
| Benchmark: change from before to during the ride | -0.136 | -0.170 to -0.101 | <.001 |
| Ride boost, companion pairs (beyond benchmark) | 0.096 | 0.050 to 0.142 | <.001 |
| Ride boost, stranger pairs (beyond benchmark) | 0.045 | -0.013 to 0.103 | .126 |
| Ride boost, stranger minus companion pairs | -0.051 | -0.106 to 0.005 | .074 |

Following synchrony through all four phases showed when the extra coupling appeared and how long it lasted (Fig 3, S6 Table). Companion pairs were slightly less synchronised than the benchmark before the ride (-0.034, 95% CI -0.064 to -0.004; P = .026). They were above it during the ride (0.062, 95% CI 0.032 to 0.092; P < .001), most clearly in early recovery (0.102, 95% CI 0.072 to 0.132; P < .001), and back at the benchmark in late recovery (0.017, 95% CI -0.013 to 0.047; P = .277). Stranger pairs looked different: they were above the benchmark in every phase, already before the ride (0.060, 95% CI 0.023 to 0.098; P = .002) and still in late recovery (0.061, 95% CI 0.023 to 0.099; P = .002). A stable gap that is present before the train leaves is more likely to reflect what riders on the same train share anyway, such as the same queue, walk and moment, than the ride itself. The time course thus adds weight to the companion result, while the stranger result looks less like ride-induced coupling.

Estimates from one linear mixed model of synchrony (Fisher z) with phase (baseline, ride), pair type and their interaction, a random intercept per pair, and the surrogate benchmark as reference. A ride boost is the change from baseline to ride in a real pair type minus the same change in the benchmark. Where riders sat, which was assigned rather than chosen, did not matter. Pairs seated side by side were no more synchronised during the ride than pairs in different cars (-0.013, 95% CI -0.109 to 0.084; P = 0.796), nor were pairs in the same car (P = 0.341), and companion pairs were somewhat less synchronised during the ride than stranger pairs once seating was taken into account (-0.054, 95% CI - 0.096 to -0.012; P = 0.012; S3 Fig), in line with the higher synchrony of stranger pairs in every phase. Riders on the same train were also far from interchangeable: the train explained almost none of the variation in heart rate (ICC = 0.007).

### Psychological change

After the ride, riders reported being happier (7.51 to 8.79; d = 0.87) and less stressed (3.46 to 2.74; d = - 0.29; Fig 4, Table 5). The largest relative change was in closeness to the other riders on the train: it more than doubled, from 1.64 to 3.84 (d = 0.70), although it stayed far below closeness to their own companions (3.83 vs. 9.00 after the ride). Closeness to companions was already near the top of the scale and rose only slightly (8.92 to 9.15; d = 0.15). All changes remained significant after correction for multiple testing. Because the upper anchor of the closeness items differed between the two surveys, the size of the closeness changes should be read with some caution.

**Fig 4.**
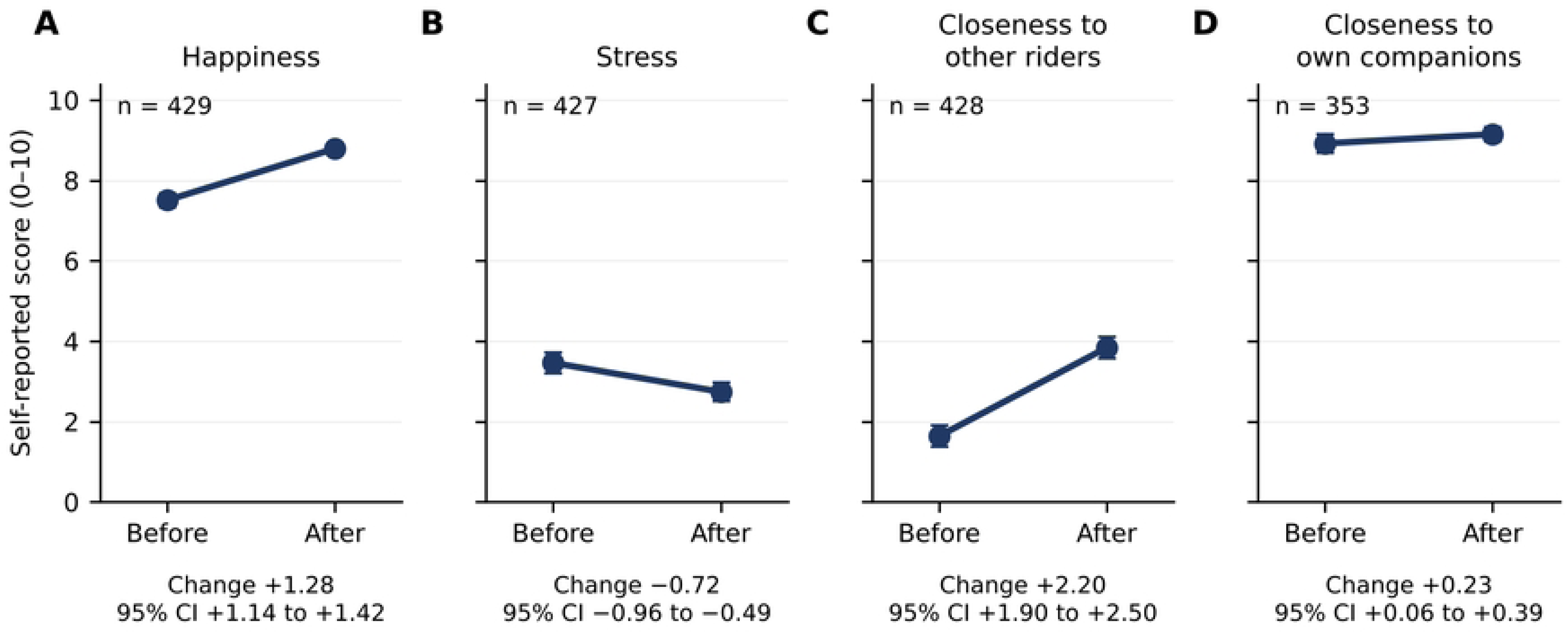
Self-reported happiness, stress and closeness before and after the ride. (A) Happiness, (B) stress, (C) closeness to the other riders on the train and (D) closeness to own companions. Dots show mean scores with 95% CIs at each survey among riders with both ratings; the change and its 95% CI from the within-person mixed model are given under each panel. All scales run from 0 to 10; higher happiness and closeness and lower stress mean a favourable change. The upper anchor of the closeness items differed between the two surveys (see Materials and methods).

**Table 5.** Self-reported happiness, stress and closeness before and after the ride.

| Outcome (0-10) | n | Before, mean (SD) | After, mean (SD) | Change (95% CI) | Cohen's d | P | q (FDR) |
|---|---|---|---|---|---|---|---|
| Happiness | 429 | 7.51 (1.76) | 8.79 (1.40) | +1.28 (1.14 to 1.42) | 0.87 | <.001 | <.001 |
| Stress | 427 | 3.46 (2.66) | 2.74 (2.47) | -0.72 (-0.96 to -0.49) | -0.29 | <.001 | <.001 |
| Closeness to own companions | 353 | 8.92 (2.21) | 9.15 (1.63) | +0.23 (0.06 to 0.39) | 0.15 | .006 | .006 |
| Closeness to other riders | 428 | 1.64 (2.90) | 3.84 (2.81) | +2.20 (1.90 to 2.50) | 0.70 | <.001 | <.001 |
| After the ride: companions vs. other riders | 375 | 3.83 (2.89) | 9.00 (1.83) | +5.17 (4.83 to 5.51) | 1.53 | <.001 | <.001 |

Changes from within-person linear mixed models. q, Benjamini-Hochberg false-discovery-rate adjusted P value across the five tests.

### Does the body predict the mind?

None of the 24 associations between phase-specific heart-rate responses and psychological outcomes survived correction for multiple testing (smallest q = 0.30; Fig 5, S5 Table). Effects were small throughout: the largest was 0.21 scale points per standard deviation of heart-rate response. Synchrony was not associated with closeness either: pairs whose heart rates moved more together during the ride did not gain more closeness to their companions (P = .208; S5 Fig).

**Fig 5.**
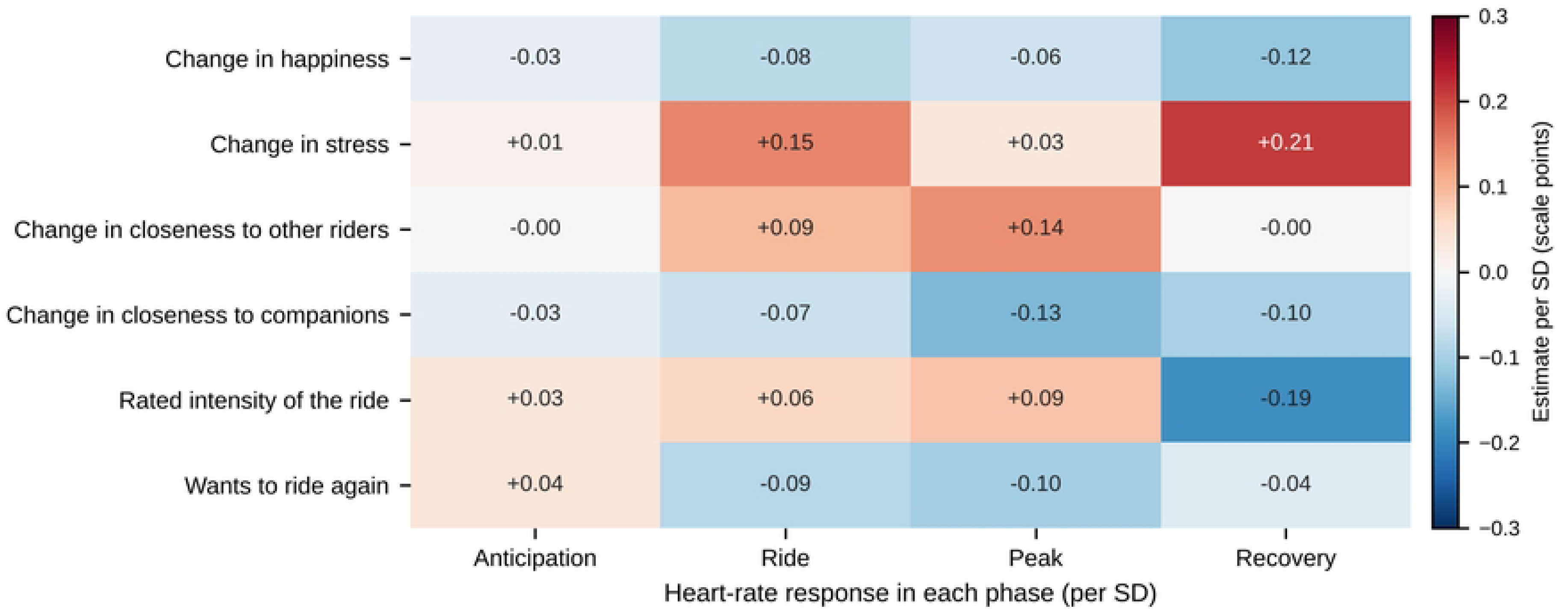
Heart-rate responses in each ride phase and psychological outcomes. Each cell shows the change in an outcome (scale points) per standard deviation larger heart-rate response in that phase, from a linear mixed model with a random intercept per train. Red, positive; blue, negative. * q < .05 after Benjamini-Hochberg correction across the 24 tests.

## Discussion

A two-minute roller-coaster ride raised riders’ heart rate by 15 bpm, with a rise that started in the last half-minute in the queue and was gone within nine minutes. Pairs of riders who had both come with companions became more synchronised than the ride alone explains; this coupling built up during the ride, peaked just after it and had faded five minutes later. After the ride, riders also reported feeling happier, less stressed and closer to the other people on the train. But how a rider’s heart responded, in any phase, was not related to how much these ratings changed.

The surrogate benchmark is what makes the synchrony result interpretable, and it behaves against intuition. Any two riders, even on different trains, were more alike in the queue than during the ride, because waiting produces a slow shared rise while the peaks of the ride hit people at different moments. Without a benchmark, one would conclude that the ride pulls riders apart. Measured against it, the ride does couple riders who came with companions. Since such riders were not necessarily together on the train, and need not have sat next to each other, this is a difference between riders who came with someone and riders who came alone, not evidence of coupling between friends. Whether it couples them more than strangers was not clear, and the answer depended on the measure, as synchrony effects often do [3]. Following the phases helps here. For companions the extra coupling appeared with the ride and disappeared afterwards, which is what a shared experience should produce. For strangers it was there before the train left and stayed at the same level, which points to the shared setting rather than to the ride. This fits the fire-walking study by Konvalinka and colleagues, where arousal was synchronised between performers and related, but not unrelated, spectators [5]. Where riders sat made no difference, unlike in everyday settings [7]; a uniform, intense stimulus may swamp the subtler effects of proximity.

We also asked whether riders fall into distinct response types. Heart-rate curves varied widely, but they did not form distinct groups. The clearest split was between riders who were calm in the queue, whose heart rate rose steeply and stayed up, and riders who were already aroused, whose heart rate rose less and then settled below their queue level. That is mainly a difference in starting point, and it was unrelated to who the riders came with, to first rides and to how the ride felt. Part of it is arithmetic: when change is measured from a single baseline value, riders who start high will tend to show smaller rises, which is why adjusting for the starting level is usually recommended [16]. In absolute terms the two profiles were about 19 bpm apart in the queue but only about 9 bpm apart at their peaks. A baseline taken in a queue is therefore itself part of the response rather than a resting value, and labels such as calm or anxious riders would read too much into the profiles. Studies that compare people on change from such a baseline should report absolute levels alongside the change and take the starting level into account.

The psychological results are substantial. Closeness to the other riders on the train more than doubled, without any interaction beyond sharing a train, although from a low level: a mean below 4 on a 0-10 scale after the ride describes a modest sense of connection, not a new relationship. One possible reason is the seating. Because seats were assigned by participant ID, riders, including those who came with companions, may often have shared the ride side by side with someone they had not come with, and sitting next to a person through an intense two minutes may be enough to feel somewhat closer to them. We could not test this, because the survey did not record who sat next to a rider’s own companions. Higher happiness and lower stress after the ride agree with field evidence that mood improves after voluntarily entering an intense, frightening setting [9]. Relief after the anticipation, having completed the ride, talking with others and the rest of the park visit may all contribute, and this design cannot separate them. Neither the heart-rate response in any phase nor synchrony was associated with these gains. This argues against a simple “more synchrony, more bonding” account and agrees with findings that physiological and felt emotion can diverge [17] and that synchrony may relate more to how groups perform than to how close they feel [6]. A null between-person association does not make physiology irrelevant, though: arousal shared by all riders could still shape how the ride feels, which comparisons between riders cannot detect. Our measure also captures coupling in overall arousal, not its timing, meaning or shared appraisal, any of which may matter more.

### Strengths and limitations

Strengths are the large field sample (439 riders on 19 trains), the surrogate benchmark, the use of each rider’s own timing within the ride, and a single scripted analysis from raw data with several robustness checks (S2 Table). The main limitations concern what the design can attribute. There was no non-riding or alternative-experience comparison group, so the psychological changes cannot be attributed to the ride itself or to sharing it; riders chose to ride and whom to come with, and most were experienced Balder riders. Seats, by contrast, were assigned by participant ID. Companion status was self-reported per rider rather than linked pair by pair, and companions were not necessarily seated together, so a companion pair may consist of two riders who came with different groups. This would tend to dilute any effect of riding with someone one knows, and we could not study pairs of actual companions. The survey did not ask whether riders knew the other people on the train, so “other riders” cannot simply be equated with strangers. The upper anchor of the closeness items changed from “very close” to “extremely close” between the surveys; a stronger top anchor would, if anything, be expected to pull post-ride ratings down, but other effects of the wording cannot be ruled out. Outcomes were single items, happiness and closeness to companions started near the top of the scale, and there was no later follow-up, so the results concern short-term states, not lasting wellbeing or relationships. Age and sex were not collected. The post-ride survey was completed several minutes after the ride, overlapping the late-recovery window, so late-phase heart rate and self-report are not strictly ordered in time. The wrist-worn optical sensor filters the signal on the device, so our synchrony measure reflects coupling over seconds, not beat-to-beat, and heart-rate variability could not be studied. The response profiles were data-driven and weakly separated, and should not be read as fixed types of riders.

## Conclusions

A short, shared thrill made the hearts of riders who came with companions beat more alike than the ride alone explains, and riders reported feeling happier and closer to the others on the train afterwards. The two ran in parallel: how strongly or how synchronously a rider’s heart responded was not related to how much closer or happier they felt. Heart-rate synchrony, at least as measured here, is not a stand-in for social bonding, and controlled comparisons are needed to tell what the ride itself adds.

## Acknowledgments

We thank Liseberg AB for operational support, the research assistants who collected the data, and all riders who took part.

## Funding, competing interests and data

***Funding:*** Liseberg AB provided partial funding and site access.

***Competing interests:*** Liseberg AB partly funded the study. The authors have no other competing interests.

## Data availability

Raw heart-rate recordings and free-text survey answers are not public because they could identify individuals; they are available from the corresponding author under a data-use agreement.

## Author contributions

Conceptualization: DB, MD. Formal analysis: DB. Writing - original draft: DB. Writing - review & editing: DB, MD, CW, MH, GW. All authors have read and approved the final version of the manuscript.

---

**Figure S1.**
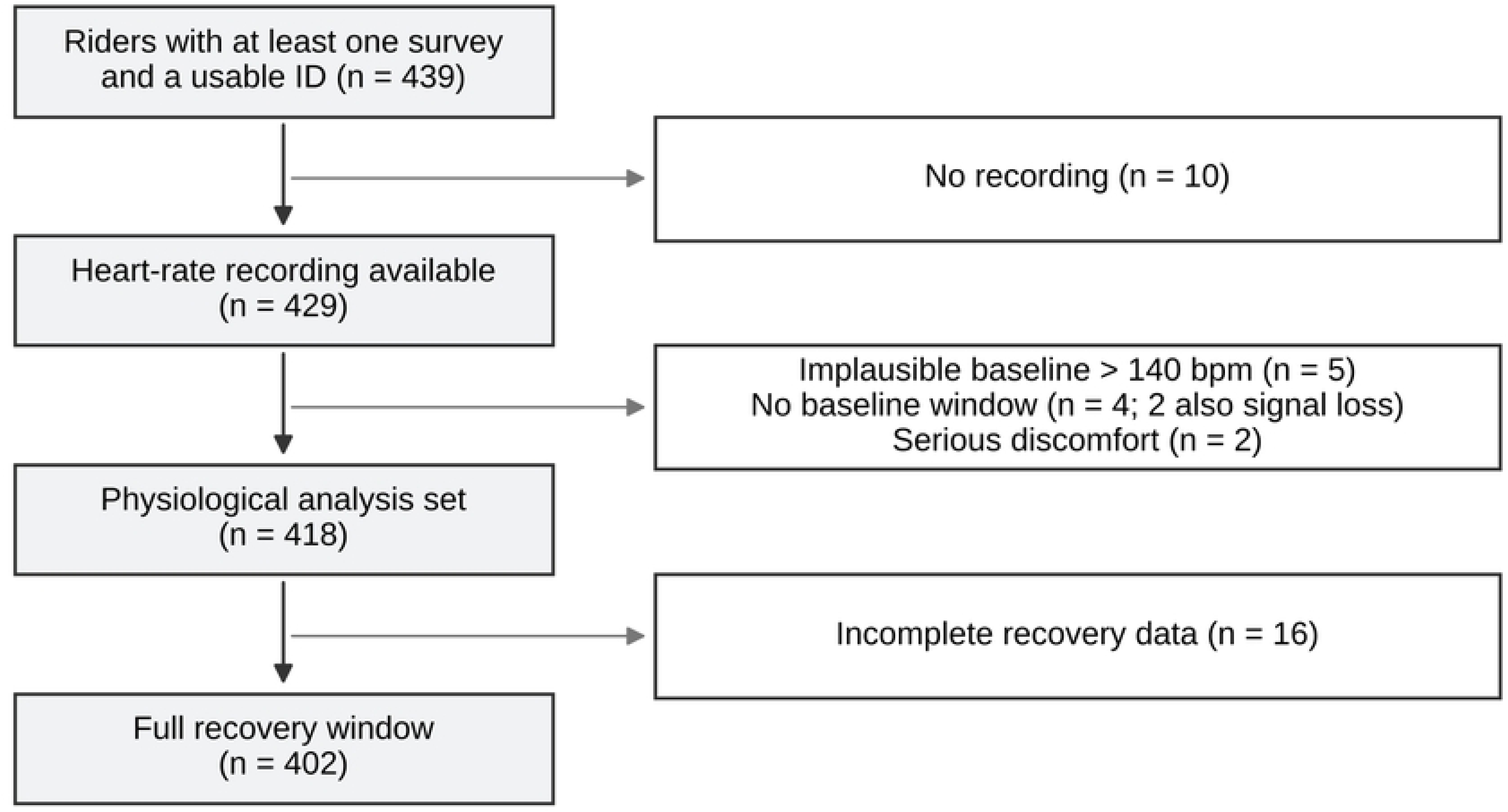

**Figure S2.**
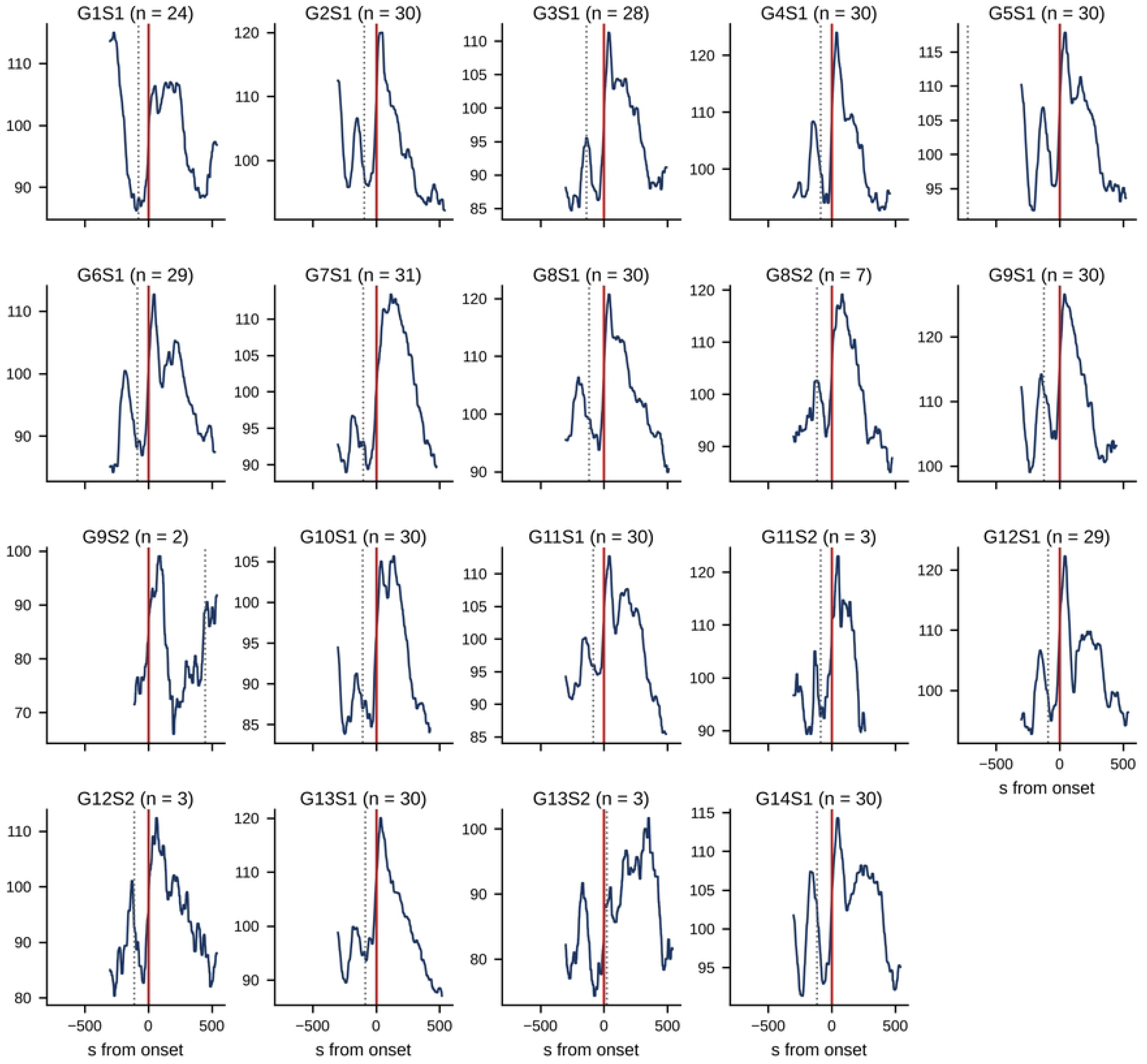

**Figure S3.**
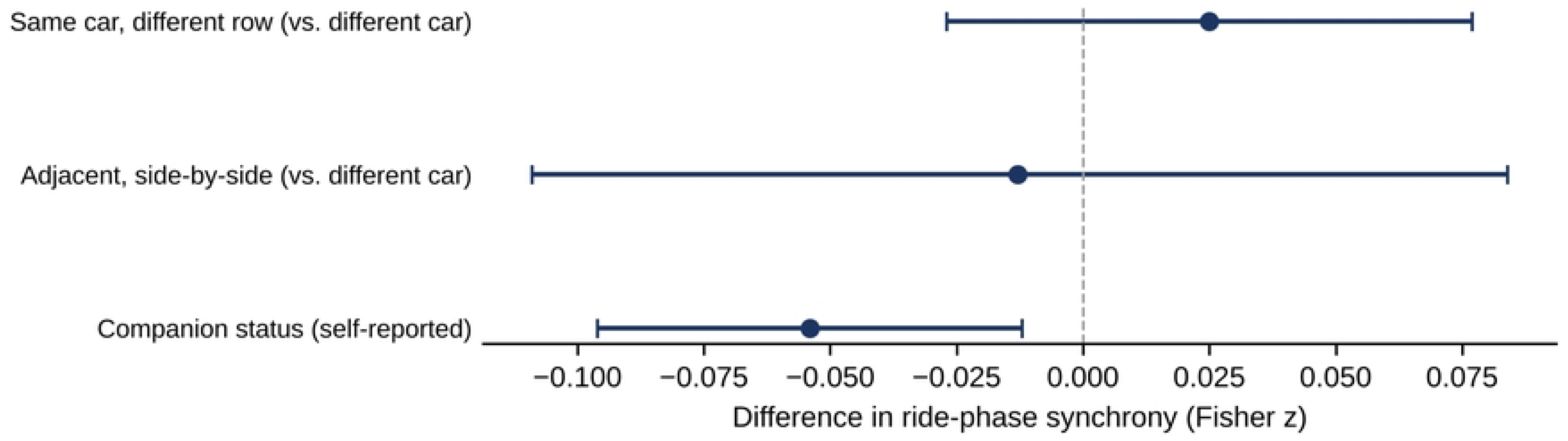

**Figure S4.**
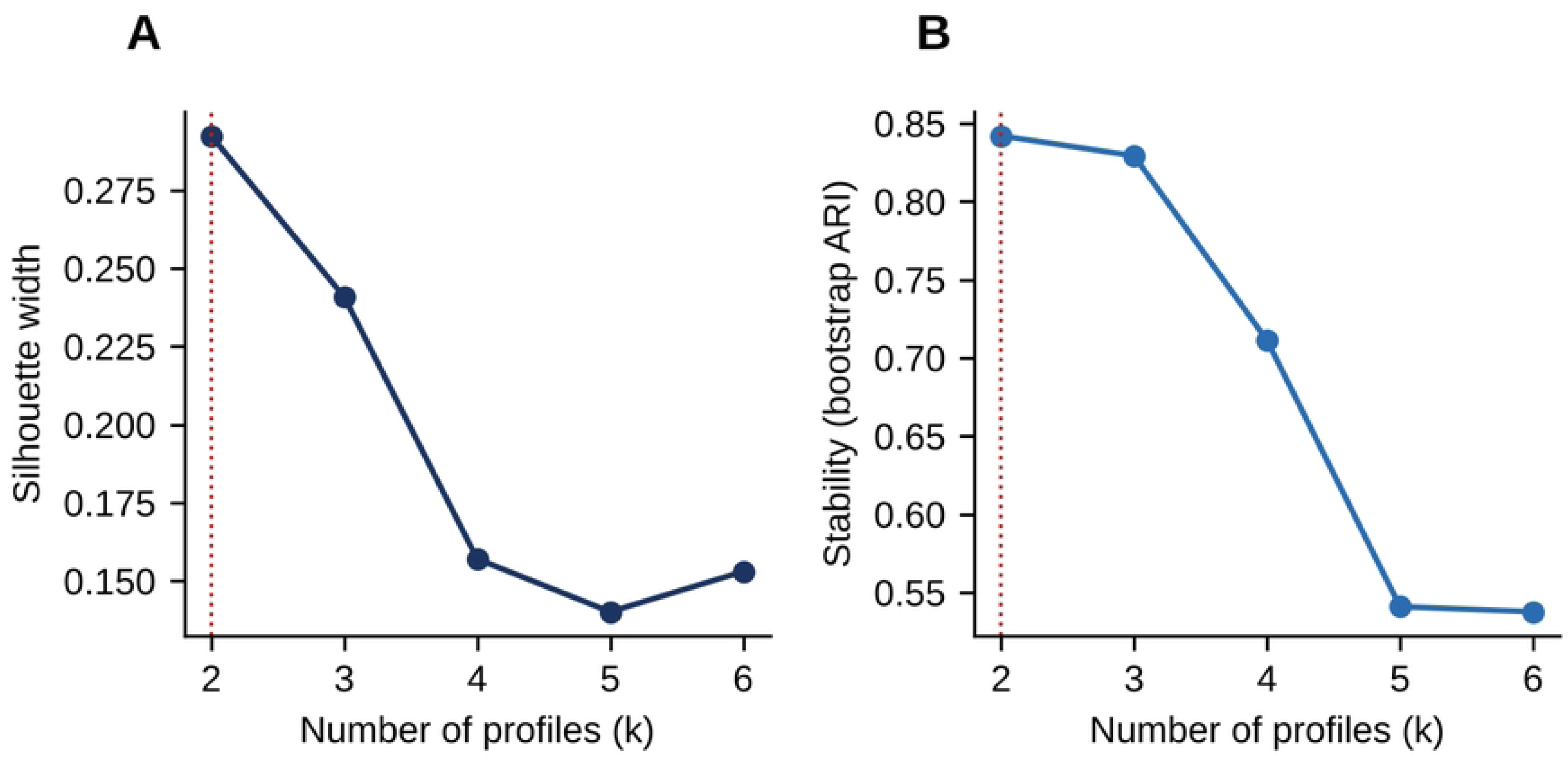

**Figure S5.**
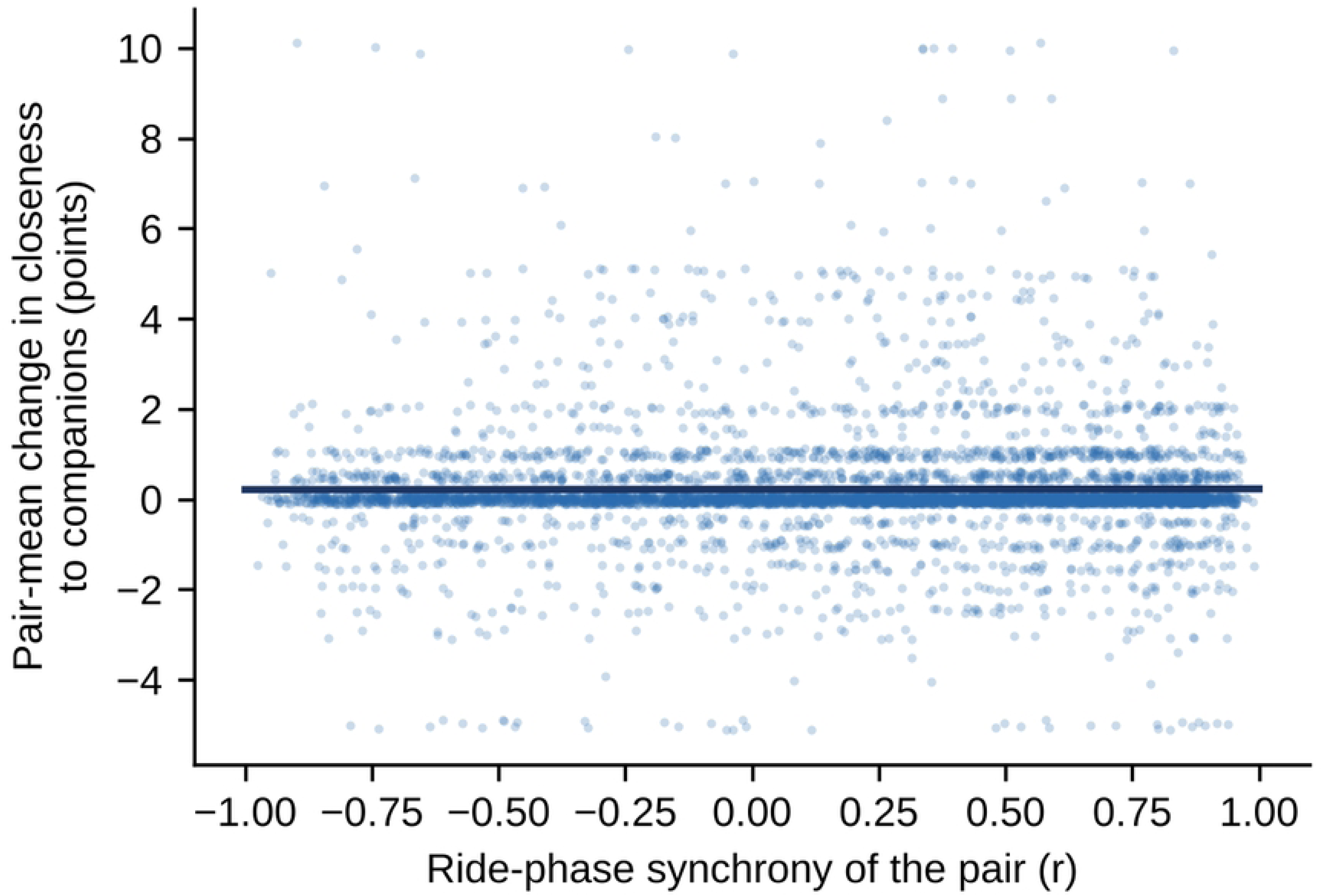

## Notes

### Competing Interest Statement

The authors have declared no competing interest.

